# Rapid plastid isolation reveals the chloroplast proteome and structures of the chlororibosome large subunit and RuBisCO in *Marchantia polymorpha*

**DOI:** 10.64898/2026.07.14.738477

**Authors:** Parth K. Raval, Cristopher Mitchell, Maria Lozano-Quiles, Sarah O’Keefe, Tuula A. Nyman, Brendan Battersby, Sarah J. Butcher, Sven B. Gould

## Abstract

Plastids house the biology of eukaryotic photosynthesis. While 1000s of plastid genomes have been sequenced, the availability of less than ten proteomes and only two species with full 70S plastid ribosomal structures limit our understanding of plant evolution. We optimized a protocol for the rapid isolation of *Marchantia polymorpha* plastids that provides a highly enriched and intact organelle fraction from gradient volumes as little as 2 mL. The approach was successfully applied to six other species. Focusing on *M. polymorpha*, we determined the proteome of the plastid fraction, identifying 1337 nuclear-encoded proteins with a high confidence, where 83% belong to orthologs shared with angiosperms. We further isolated protein complexes by RNA affinity purification using poly-lysine and provide the high-resolution structures of the 50S subunit of the chloroplast ribosome and RuBisCO using cryogenic EM and image reconstruction to 2.23 and 2.12 Å resolution, respectively. For plastids, our data show that the genome reduction event experienced by the bryophyte common ancestor has had little impact on the organelle’s complexity and they underscore a high level of structural conservation of key components. Our data provide novel resources to explore the functional evolution of plastid proteomes and major macromolecular complexes of cyanobacterial origin.

## Introduction

The transition from a cyanobacterial endosymbiont to the different types of plastids of algae and plants entails the loss and transfer of most of the endosymbiont’s genetic information^1–5^. Crop plants such as pea or maize hence encode more than a thousand proteins in their nuclear genome that need to be imported by their plastids after cytosolic translation^6,7^. Plastid proteomes are complemented by a few proteins that are kept organelle-encoded, maybe due to the need for the co-localisation of genes of the electron transport chain and the bioenergetic membrane itself for functional redox regulation^8–10^, and protein hydrophobicity impairing membrane translocation^11–14^. Subunits required during the early steps of 70S ribosome assembly are likely retained due to constraints associated with securing proper ribosome maturation^15^. Typically, as is the case for the 70S ribosome, plastid-encoded proteins interact with nuclear-encoded ones to form macromolecular complexes such as photosystem I and II, the NADH dehydrogenase complex, the ribulose-1,5-bisphosphate-carboxylase/-oxygenase (RuBisCO) enzyme, and ATP synthase.

A significant event in plastid evolution was the transition from water to land^16,17^. Plant terrestrialization and the origin of the Embryophyta is characterised by the emergence of the cuticle, stomata, spores and seeds, multiplastidy, the adaptation of stress response networks already present in the streptopyhte algal ancestor, a step-wise expansion of genomes, and a general increase in mitochondrial and plastid proteome size^17–23^. An early-branching and ecologically important group of land plants are the bryophytes, uniting liverworts, hornworts, and mosses (Fig. 1). After genome expansions at the origin of the Chloroplastida and again at the origin of Streptophyta^24,25^, the bryophyte ancestor experienced genome reduction that drove the lineage’s phenotypic evolution^19^. Now, after more than 500 million years, bryophytes are characterized by highly derived genomes that in sum encode a larger variety of gene families than their vascular relatives^26^.

**Figure 1:**
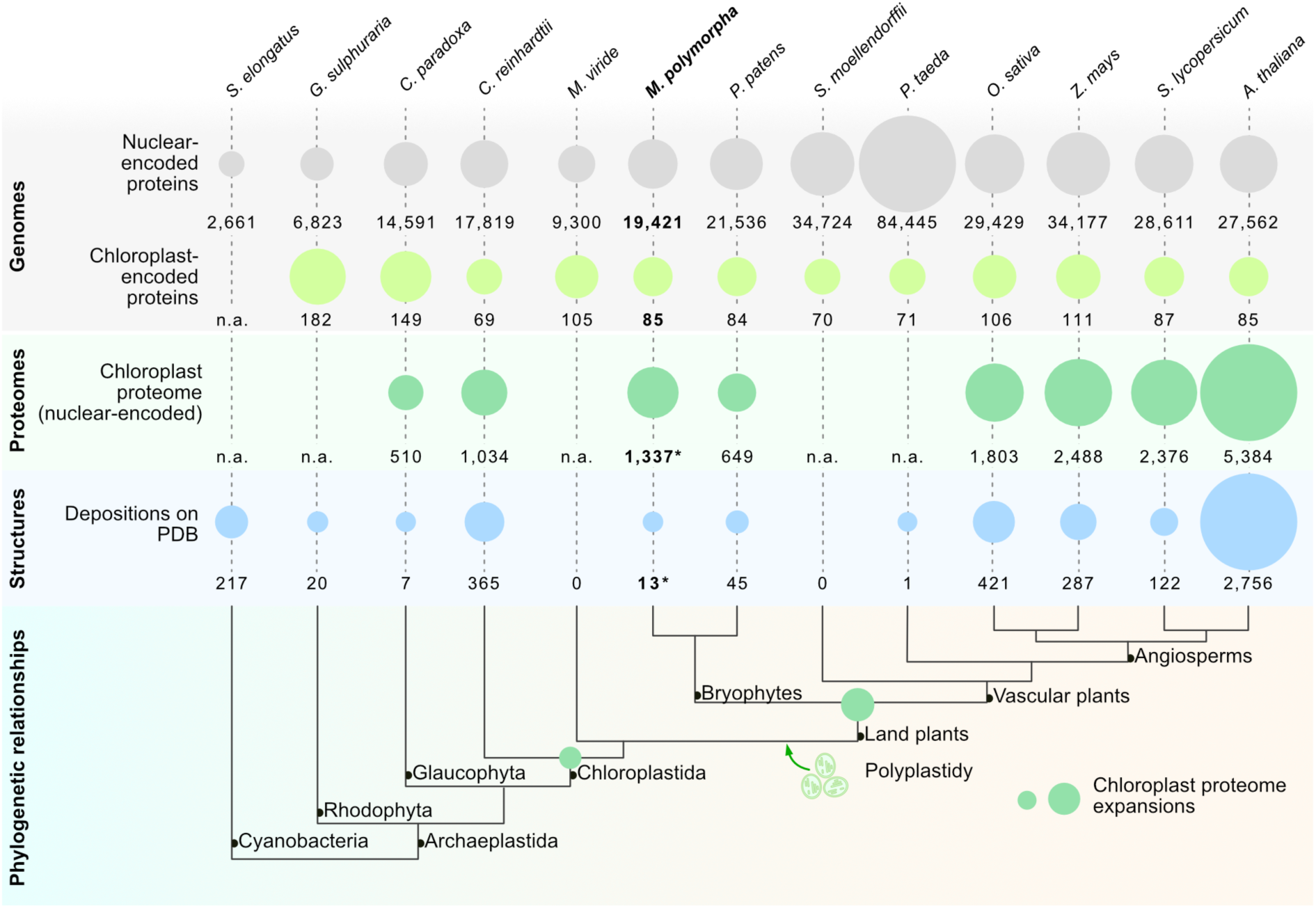
Phylogenetic relationships and key numbers for exemplary species. A cladogram of key species representing the diversity of Archaeplastida lineage, with the number of nuclear- and chloroplast-encoded proteins and the number of proteins in the chloroplast proteome and the number of entries in the World-Wide Protein Data Bank (last accessed on 1^st^ May 2026). Asterisks indicate numbers that also include the current study. Key changes in plastid biology over the course of plant evolution are indicated on the cladogram^18,20,28,29^. The number of nuclear encoded proteins as per the whole genome models^30^ and the number of plastid localized proteins from previous studies: *Cyanophora paradoxa*^31^*, C. reinhardtii*^32,33^, *Physcomitrium patens*^34^, *Oryza sativa*^35^, *Zea mays*^35^, *Solanum lycopersicum*^35^, and *A. thaliana*^36^ (N.a. = not available; species from the left: *Synechococcus elongatus, Galdieria sulphuraria, Cyanophora paradoxa, Chlamydomonas reinhardtii, Micromonas viride, Marchantia polymorpha, Physcomitrium patens, Selaginella moellendorffii, Pinus taeda, Oryza sativa, Zea mays, Solanum lycopersicum, Arabidopsis thaliana*).

The positioning of bryophytes on the phylogenetic tree provides an ideal opportunity to explore the trajectory of plastid biology after genome reduction in comparison to vascular plant plastids. Experimentally curated plastid proteomes and high-resolution structures of macromolecular complexes outside of a few angiosperm model systems, however, are scarce (Fig. 1). To address this, we optimized a chloroplast isolation protocol that requires minimal resources and time. It yields intact chloroplasts from a diverse set of species and eases their study, enabling chloroplast biology exploration across a broad variety of taxa. Focusing on the liverwort *Marchantia polymorpha,* we obtained a highly enriched chloroplast proteome, which corroborates previously predicted evolutionary trajectories of land plant plastid proteomes^20^. With the aim of isolating chlororibosomes, we subjected the plastid fraction to an RNA affinity purification using poly-lysine^27^. This resulted in the enrichment of multiple macromolecular complexes from the isolated chloroplasts. Using cryogenic electron microscopy (cryo-EM), we resolved the structures of the 50S ribosomal subunit and RuBisCO at respective resolutions of 2.23 and 2.12 Å. Our data provide new resources to study plastid biology and allow us to hypothesize that the chloroplast proteome and ribosome of the bryophyte were not significantly affected by genome reformatting at the origin of the bryophytes.

## Results

### A rapid mini-prep protocol for the isolation of plastids from diverse species

Plastid isolation protocols are available for very few species and they can be complex and resource intensive. A widely applicable and efficient purification protocol has the potential to accelerate and expand plastid research across diverse plant lineages. To this end, we systematically scaled down a Percoll^®^ gradient-based isolation protocol and tested it first on the bryophyte *Marchantia polymorpha*. From as little as 500 mg of starting tissue material (i.e. only a few, two-week old thalli), we observed the formation of a characteristic dark green band at the Percoll^®^ interface, which was confirmed as clean and intact chloroplasts by microscopy (Fig. 2a). We tested our protocol on standard model systems such as *Arabidopsis thaliana* and *Nicotiana tabacum*, as well as *Moricandia arvensis* (a model for C3-C4 intermediate photosynthesis), *Diplotaxis viminea* (a model for stress physiology), and *Erucastrum gallicum* (a model in plant ecology). Our protocol performed equally well across each model system, taking under one hour to complete, from less than 1 g of leaf tissue for each gradient (Fig. 2a). For *Chlamydomonas reinhardtii*, we scaled down a previously reported protocol requiring a three-, rather than a two-, step Percoll gradient^37^ and found it to be similarly effective for the algal model system (Fig. 2a).

**Figure 2:**
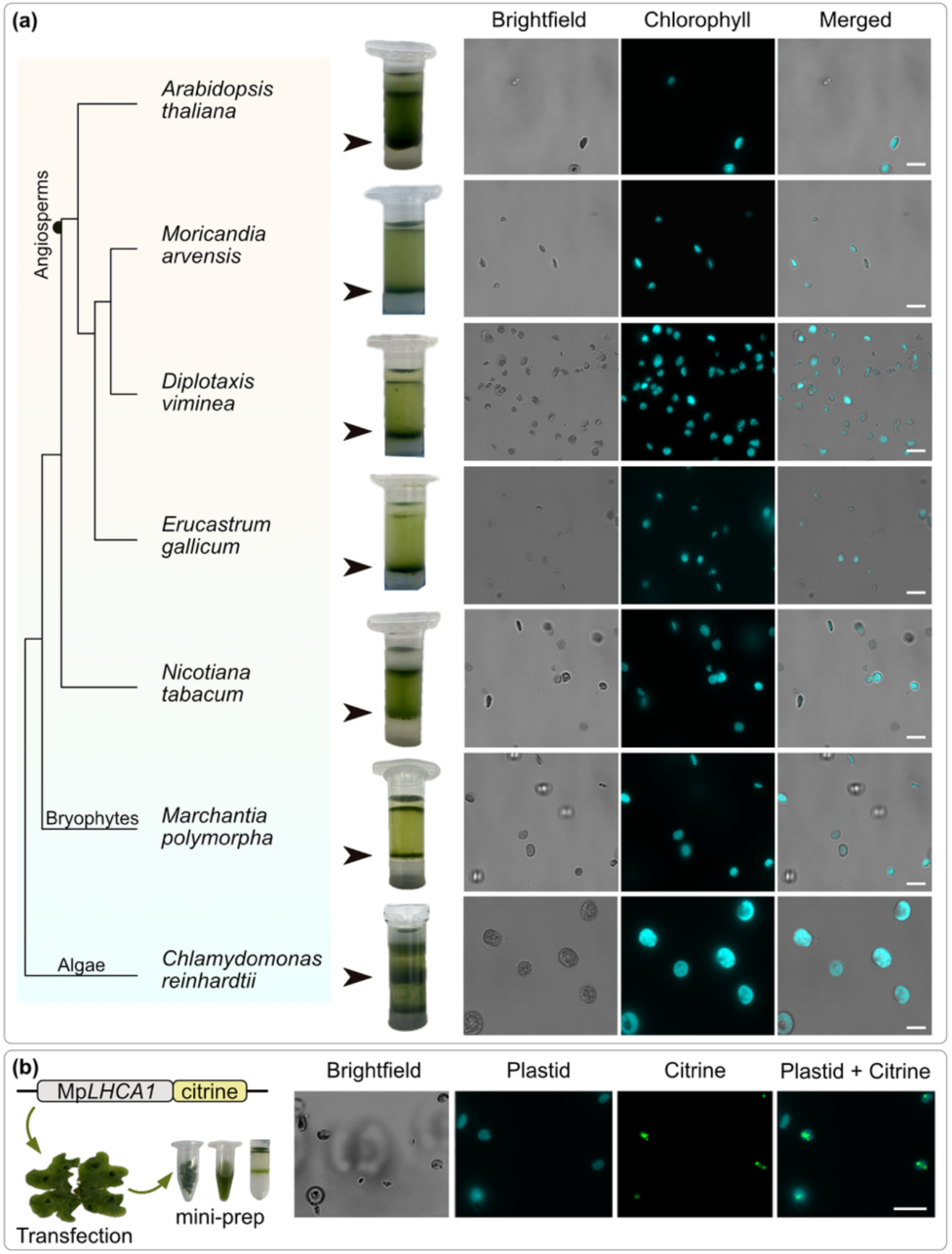
Small-scale plastid isolation from diverse species. **(a)** Plastids isolated from diverse species using the mini-prep protocol and visualized through their chlorophyll autofluorescence. Scalebars 10µm. **(b)** Chloroplasts isolated from a *M. polymorpha* transfectant line expressing LHCA1:citrine shows colocalization of chloroplasts and the citrine signal.

Subcellular fractionation is the first step in characterizing the proteome of a compartment. After the isolation of the *M*. *polymorpha* chloroplasts, we assessed the fraction microscopically alongside a transfectant line expressing the citrine-tagged light-harvesting complex A1 protein (LHCA1; Mp2g15420). This showed a clear colocalization of the reporter with autofluorescence and the shape of the purified organelles (Fig. 2b). We further confirmed the purity of the isolated chloroplast fraction by Western blotting using antibodies specific for established marker proteins of other compartments, such as BiP1 for the endoplasmic reticulum and SHMT for mitochondria (Fig. S1). From biological triplicates of chloroplasts isolated from individually grown plants, tandem mass spectrometry (MS) identified 1337 nuclear- and 43 plastid genome-encoded (Table S1). 61.7% of the nuclear encoded proteins identify a homologue in at least one of the 56 diverse cyanobacteria that were part of our protein database for screening (Fig. S2), thus underscoring the high degree of proteome conservation of this photosynthetic organelle of eukaryotes over a span of more than 1.5 billion years^38^.

A functional annotation of the chloroplast proteome identified here shows that most proteins play a direct role in photosynthesis, with functions related to chlorophylls, protein synthesis, carbon and amino acid metabolism and chloroplast-specific secondary metabolism pathways such as cofactor and pigment biosynthesis (Fig. 3a). These categories, expected of a chloroplast proteome, speak for the purity of the isolated fraction. The largest single fraction of proteins (184 proteins) returned no clear annotated function in the MarpolBase^39^ (accessed 1^st^ August 2025). 125 of these 184 proteins had homologues in *A. thaliana* (Table S2A), of which 31 are involved in thylakoid and ribosome biogenesis, RNA processing and localized in thylakoid membranes, but the remaining 94 homologues had no clear functional annotation for Arabidopsis either. 38 of these 184 proteins lack homologs in angiosperms and three appear unique to *M. polymorpha*. A screen for known domains identified 13 proteins encoded tetratricopeptide repeats (TPR), two encoded pentatricopeptide repeats (PPR) and 20 encoded RNA related domains, making TPR/PPR and RNA related proteins the single largest fraction among the 184 proteins with an uncertain function (Table S2B). For 13 proteins (Table S2C), however, no functional domains could be predicted but we found a match for two of them by predicting the structures with AlphaFold3 and using Dali^40,41^ to search for structural homologues (Mp3g01370 aligned with PsbQ and Mp5g18410 with an AFMP4P virulence factor), leaving eleven plastid proteins that require experimental investigation.

**Figure. 3:**
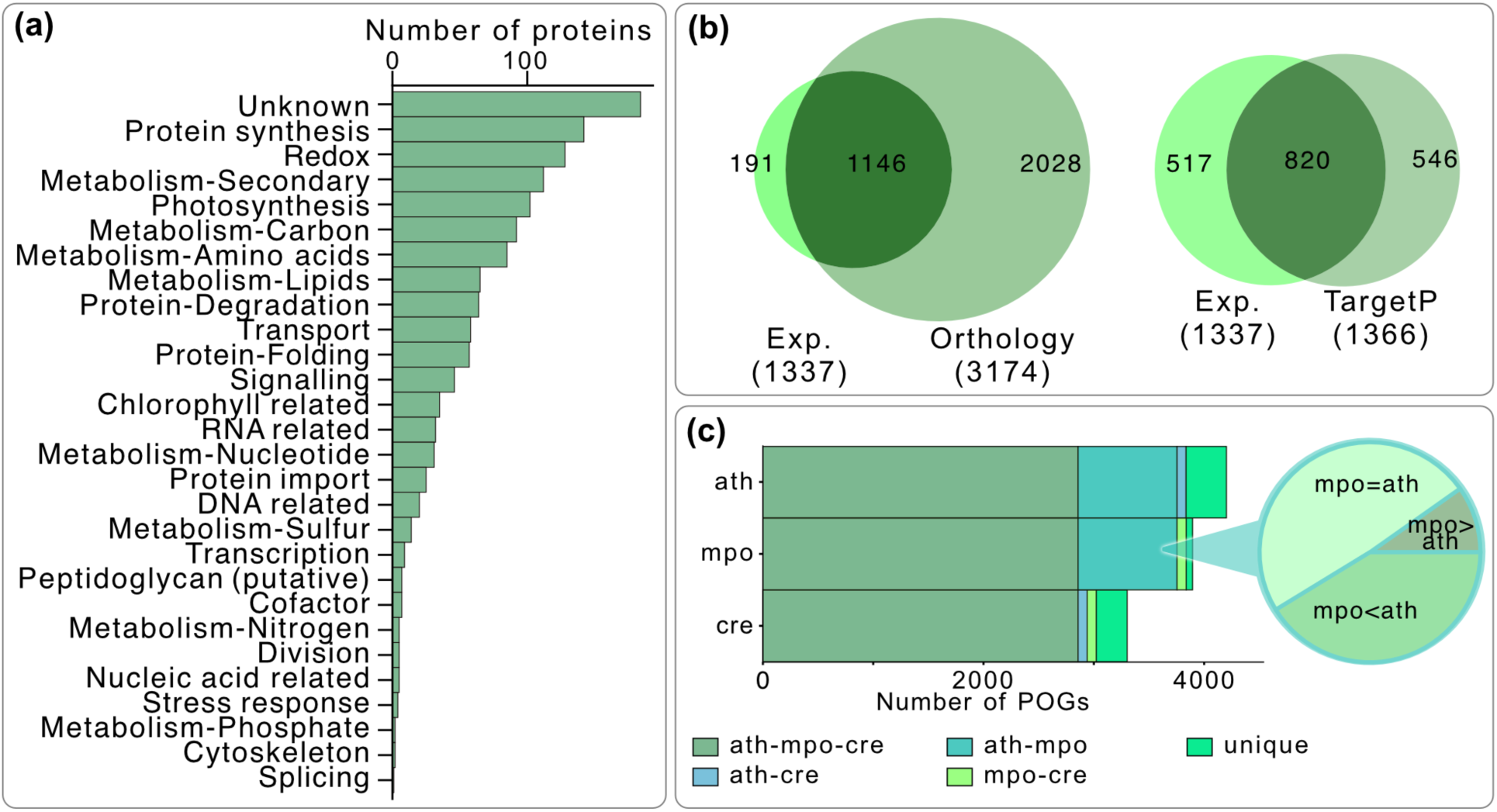
The *M. polymorpha* plastid proteome gained almost as many proteins since genome remodeling at the origin of bryophytes as it had lost. (a) Distribution of the identified plastid proteins across functional categories. **(b)** Comparison of the experimentally-curated Marchantia plastid proteome with bioinformatically predicted orthology groups^50^, and proteins predicted to be plastid targeted by TargetP v2.0 **(c)** Comparisons of POGs of *A. thaliana* (ath), *M. polymorpha* (mpo) and *C. reinhardtii* (cre), with each stacked region representing the number of shared POGs. Of the 897 POGs shared between *A. thaliana* and *M. polymorpha*, half are represented by a similar number of proteins per group, while about 40% are reduced and 10% are expanded in the number of genes in *M. polymorpha* in comparison to *A. thaliana* (pie chart on the right).

### Lineage-specific gene family expansions contribute to the *M. polymorpha* plastid proteome

Comparative proteomics is an important component in understanding organelle evolution and the contribution of the compartment to the ecophysiology of a plant. Ancestral state reconstruction of mitochondrial and plastid proteomes across a billion years of chloroplastidal evolution recently underscored the expansion of organelle proteomes from algae to angiosperms^20^. It predicted an intermediate chloroplast proteome size for *M. polymorpha* and we find that 85.6% of the experimentally-curated nuclear encoded chloroplast proteins were predicted based on orthology with previously experimentally characterized proteins (Fig. 3b). Apart from corroborating the identity of individual chloroplast proteins, it also shows that an orthology-based approach to predict an organelle proteome has merit for plotting broader evolutionary patterns in the absence of an experimentally-curated proteome. Hence, we clustered the genomes of 57 diverse chloroplastidal species (Table S3) into orthogroups following the approach used previously^20^ and focused on 4,797 plastid orthogroups (POGs). The comparisons of the POGs and of the experimental proteome recapitulates the expansion of the chloroplast proteome from algae to angiosperm^20^, with that of *M. polymorpha* – by sheer number of POGs – ranging in the middle (Fig. 1, Fig. S3).

For 2857 POGs, a representative protein has been experimentally localized to the plastid of at least *M. polymorpha*, *A. thaliana*, or *C. reinhardtii* (Fig. 3c). 897 POGs are shared by both *A. thaliana* and *M. polymorpha*, but no protein from the *C. reinhardtii* genome was present in these POGs (Fig. 3c), suggesting they were likely gained or assigned to the chloroplasts of the Embryophyta ancestor (e.g. after duplications) and retained by Marchantia. The latter would have happened in the background of a global genome reduction event postulated for the bryophyte ancestor^19^. 48.9% of these POGs contain fewer gene members in *M. polymorpha* than in *A. thaliana*, as expected from ancestral genome reduction. 41.2% of POGs, however, contain the same number of genes in *A. thaliana* and *M. polymorpha* and 9.8% (88 POGs; largely associated with secondary metabolism and redox balance) contain more genes in *M. polymorpha* (Fig. 3c), the latter hinting at lineage-specific expansions. Comparisons of POGs across 57 species also identifies 341 POGs that are absent in more than 90% of vascular plants and present only in algae or bryophytes (Table S3). 91% of these POGs remain uncharacterized and they likely contain lineage-specific expansions that helped bryophytes to adapt to a terrestrial lifestyle in ways that differ from tracheophytes.

Even for those categories that are not uniquely expanded, such as those of translation, redox balance or detoxification (Fig. 3a), their evolution is informative. For instance, in angiosperms the NDH super complex is stabilized by PSI linkers (i.e. LHCA5 and LHCA6). This NDH-PSI interaction is crucial for photoprotection and cyclic electron flow under diverse light conditions encountered by land plants^42–45^. Based on the consistent distribution of the linker subunits across angiosperms, as compared to their sporadic presence outside of angiosperms, a stepwise evolutionary acquisition of these linkers has been proposed^46–49^. We identified peptides for the baseline subunits of the NDH complex such as subcomplex A and E, and the core LHCA1-4 subunits, while no linkers were detected (Table S1). As such, our observation supports the view that a monomeric NDH complex preceded the NDH-PSI super complex. Future structural studies on the liverwort’s NDH complex and an interpretation in light of secondary genome reduction in bryophytes is hence key in judging the credibility of a monomer first model.

### The structures of the 50S ribosomal subunit and RuBisCO demonstrate a high level of macromolecular complex conservation

With the aim of determining a chlororibosome structure for *M. polymorpha*, we subjected the purified organelles to RNA affinity purification by poly-lysine (RAPPL)^27^ . The eluate of this purification was used for mass spectrometry, which showed an enrichment of all chlororibosome subunits, the ribulose-1,5-bisphosphate carboxylase/oxygenase (RuBisCO) enzyme, and proteins of PSII, PSI, and the ATP synthase (Table S4). The eluate was used for cryo-EM grid preparation and subsequent single-particle reconstruction. 2D classification revealed several large protein complexes including recognizable views of the chlororibosomal large 50S subunit (cLSU), RuBisCO, the soluble F1-ATPase complex, and a chaperone (Fig. S4). Of these, there were sufficient varied views of RuBisCO and the 50S for three-dimensional reconstruction. No particle classes containing the full 70S or the small 30S ribosomal subunit were identified.

We reconstructed high-resolution density maps of the cLSU and RuBisCO to respective resolutions of 2.23 and 2.12 Å, reported at the 0.143 Fourier shell correlation cut-off^51^ (Fig. 4a & 5a, S4-5). These maps were used for atomic modeling using previously published sequences^52–58^. The ribosomal protein (RP) structures were first predicted using AlphaFold3^41^ and then fitted and refined into the experimental map. The rRNA was mutated from a *Spinacia oleracea* model (PDB ID 5MLC^59^). The quality of the density map allowed for modeling 6 rRNA modifications in the cLSU 23S rRNA chain (Fig. S6). Out of the 6 modifications, 5 were identified previously in Chlamydomonas and *Escherichia coli* structures, whereas the remaining modification, U579 to 5-methyluridine, appears to be novel to Marchantia (Fig. S6)^60–62^. Alignment of the spinach 50S to the Marchantia 50S revealed only 5 subunits out of 31 with a root mean square deviation (RMSD) above 3.0 Å. These were L5, L13, L21, L33, and PSRP6, with RMSDs of 3.7, 8.1, 3.1, 3.2, and 4.5 Å respectively. The values reported for L5 and L33 are due to variation in the position of a single small loop region in each. The main difference between PSRP6’s across the structures is that 9 more C-terminal residues were modeled in the Marchantia model, however these residues are also present in the spinach PSRP6 sequence, so this represents a difference in flexibility or map quality. The main noticeable difference between the Marchantia and spinach models was found at the surface near the exit tunnel, where RPs L13, L21, and L22 are present (Fig. 4b). In *S. oleracea*, these RPs have regions at their N- and C-termini that stick out from the surface and they were hypothesized to form an interface with the thylakoid membrane^59^.

**Figure 4:**
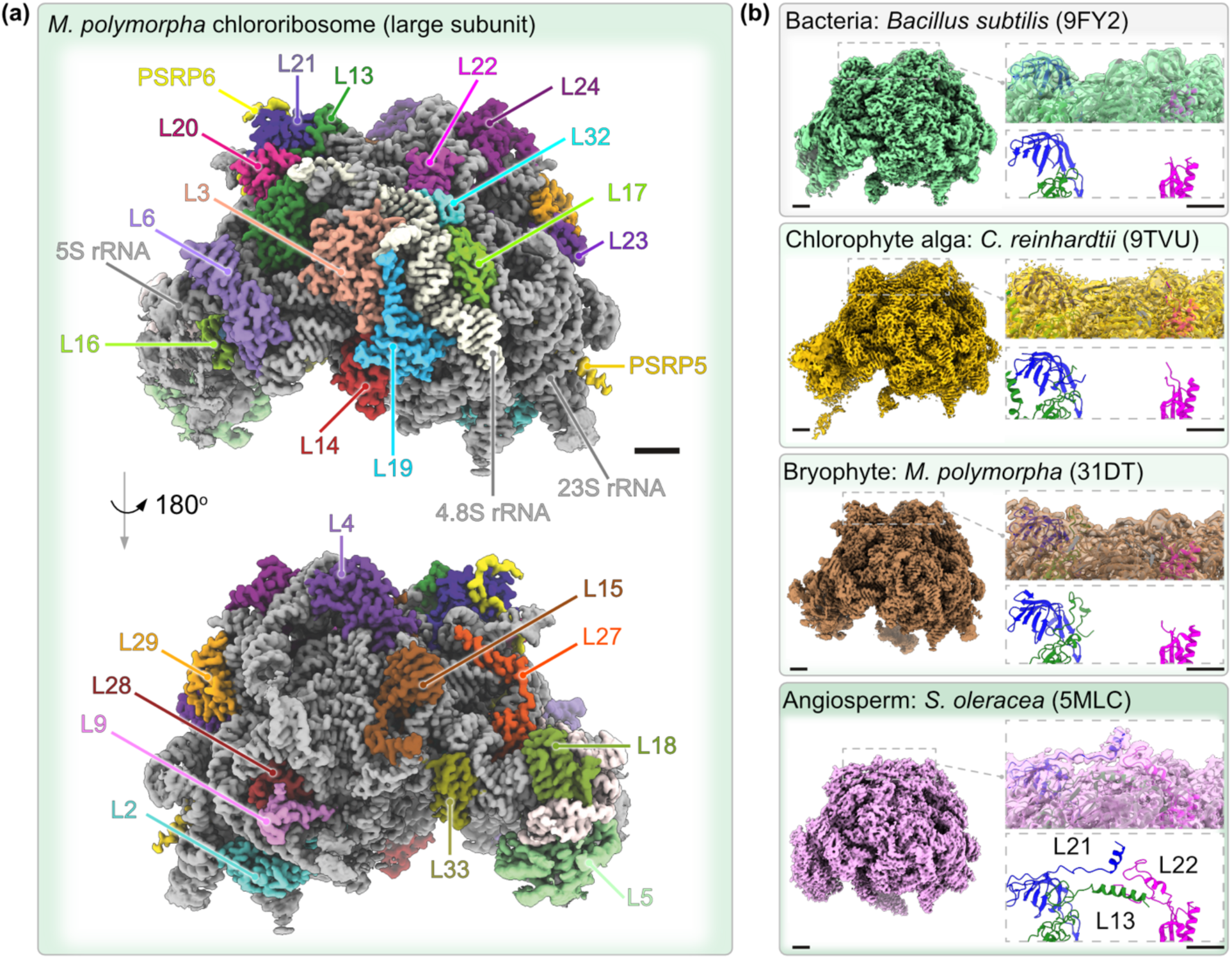
The *M. polymorpha* chlororibosome 50S subunit differs from that of spinach in a region proposed to interact with the thylakoid membrane. (a) Sharpened map of Marchantia cLSU, zoned around model and colored according to individual RPs and rRNAs shown as a surface at 1 σ above the mean. **(b)** Comparative density maps (depicted as surfaces) of the LSU’s from the bacterium *Bacillus subtillis* (PDB ID 9FY2, zoned around LSU model, 3 σ above the mean), chlororibosome from the alga *C. reinhardtii* (PDB ID 9TVU, zoned around LSU model, 6 σ above the mean), the angiosperm *S. oleracea* (PDB ID 5MLC, 2 σ above the mean), and *M. polymorpha* chlororibosome (PDB ID 31DT, 1 σ above the mean). Images on the right show atomic models in cartoon representation focused on proteins L13 (green), L21 (blue), and L22 (magenta) with density maps depicted as transparent surfaces at thresholds of 3, 4, 2, and 1 σ above the mean (respective to order depicted, top to bottom) and surrounding RPs and rRNA colored gray. Scale bars 20 Å.

Comparisons of structure and sequence suggest that the C-terminal region was extended (e.g. L21) or gained (e.g. in L22) by the ancestor of eudicots, since they are absent or shorter in other chloroplastidal species, including *M. polymorpha* (Fig. 4b). The N-terminal sequence of L13 is present in the annotation of the Marchantia protein but does not appear in the density map (Fig. 4b). Overall, while the amino acid sequences for chloroplastic RPs between the *S. oleracea* and *M. polymorpha* 50S share on average 68% identity, they exhibit highly similar structures through conserved folds (Table S5).

RuBisCO is well known for its high level of structural conservation^63^ and the structure of the *M. polymorpha* enzyme makes no exception. It aligns well with structures of different species spanning more than a 1.5 billion years of diversification that we compared it to (PDB ID 9FWV, 1IWA, 1GK8, 9N37, 4RUB, 9CQ5, 4HHH^64–71^). This is evident in the visual comparison (Fig. 5), and the z-scores calculated in Dali^40^ where a z-score above 8 indicates likely homology and a score above 20 indicates definite homology. The z-score is influenced by matching secondary structural elements so, the *M. polymorpha* large and small subunit chains were separately compared in Dali. For the chains of the large subunit, which include the enzyme’s active site, the z-scores ranged from 53.4 for *T. antarctica* to 57.5 for *N. tabacum* with a maximum possible score of 66.2 (Fig. S7). For the small subunit, which shows some variance in a single loop region (D176-T198), the z-scores ranged from 15.1 for *G. partita* to 19.9 for *S. oleracea* in comparison to the maximum possible score of 24.5 (Fig. S7).

**Figure 5:**
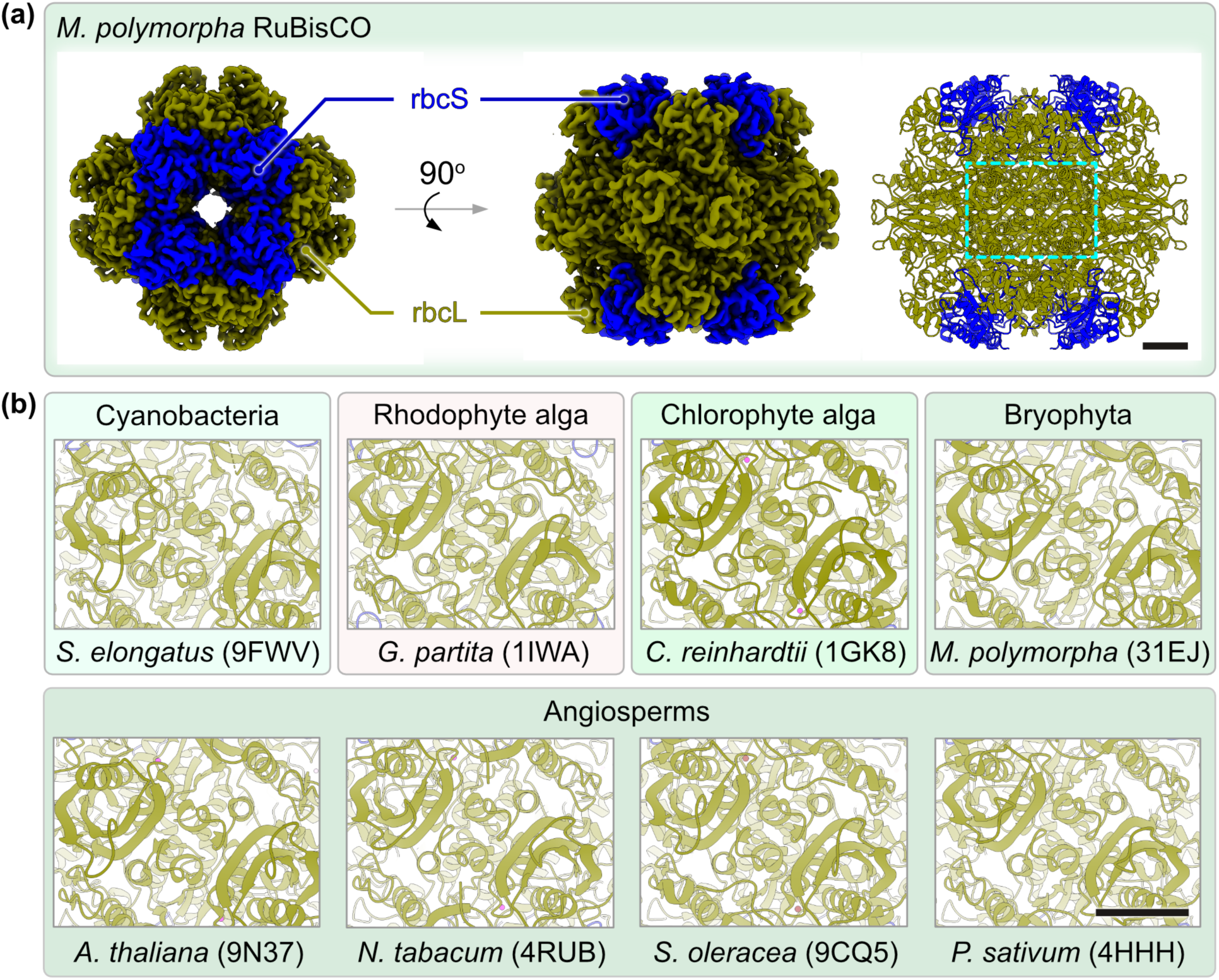
The RuBisCO structure of *M. polymorpha* underscores the enzyme’s high level of conservation since the origin of the hetero-oligomeric form I in cyanobacteria. **(a)** 2.12 Å density map and atomic model of *M. polymorpha* RuBisCO with the small subunits shown in blue and the large subunits shown in green, depicted as a surface, colored by the model at a threshold of 3 σ above the mean; cartoon model on the right with a region containing 2 active sites in adjacent large chains highlighted by a dashed rectangle. **(b)** An assembly of the same region as highlighted in (a) of eight resolved RuBisCO structures from diverse photosynthetic eukaryotes and a cyanobacterium to highlight the level of conservation spanning >1.8 billion years of evolution. PDB IDs are provided next to the species name. Ions are present in the active sites in 4 of the structures (Mg^2+^ colored pink and Mn^2+^ colored red). Scale bars 20 Å.

## Discussion

Here, we have developed a rapid and efficient plastid purification protocol that is effective across six tested plant lineages. By combining our plastid isolation approach with a RAPPL protocol to enrich chloroplast protein complexes, we provide high-resolution structures of the large subunit of the plastid ribosome and RuBisCO from *M. polymorpha*. Together with the plastid proteome, our high-resolution structures of two key protein complexes present a unique window into land plant evolution, as bryophytes are characterized by an ancient genome reduction event and a subsequent gene family expansion independent from that of vascular plants^19,26,72^.

The oldest fossil record of a plastid is that of the filamentous red alga *Rafatazmia chitrakootensis*. Their rhodoplasts were captured in 1.6-billion-year-old rock formations of the Tirohan Dolomites from central India^38^. At that point in time, these red algae (rhodophyte) plastids had already evolutionarily diverged from those of their common ancestor of Archaeplastida, from which the Glaucophyta and Chloroplastida (the green lineage) also evolved^73^ (Fig. 1). Ever since, the photosynthetic organelles of algae and plants have experienced constant adaptations, with the transition to a terrestrial habitat unequivocally representing a groundbreaking step for the evolution of life.

Bryophytes occupy an evaluative position in the evolution of land plants. They are the sister-clade to all vascular land plants^72,74^, have retained ancestral features such as flagellated sperms or a dominant, photosynthetic haploid gametophyte^75–77^, and their genomes have independently diverged from those of vascular plants after having been reduced in their common ancestor^19,26^. We find the Marchantia chloroplast proteome to comprise just over 1300 proteins, of which more than 80% are represented by orthologs in angiosperms. The large number of orthologs speaks for a trade-off between ancient nuclear genome reduction and the need to retain – or even expand – gene families needed for the complexity of plastid biology that had to adapt to a terrestrial habitat. We predict that additional organelle proteomes from bryophytes will support this pattern.

Annotating the *M. polymorpha* plastid proteome finds almost 200 proteins of unknown function, with only sporadic domains being identifiable in some cases. TPR and PPR proteins were particularly enriched. TPR domains are key for protein-protein interactions and are involved in chloroplast chlorophyl biogenesis, photosystem assembly and intracellular chloroplast distribution^78–81^. PPR domains are crucial for plastid RNA editing and organelle-targeted members of this protein family expanded with the origin and diversification of land plants, probably to compensate for an increase in mutations caused by higher levels of UV radiation ^20,82–85^. Theory has it that complex thalloid liverworts such as *M. polymorpha* lack mRNA-editing possibly due to secondary loss^86,87^, but recent results on *Cyathodium cavernum* revealed a minimum of 129 canonical C-to-U mRNA editing sites, suggesting that this is not a universal trait of the taxon^88^. Studies thus far describe on average 100 proteins with PPR (or related RNA processing) domains encoded by liverwort genomes^89,90^, though their intracellular localization was not clear. We find 35 proteins (including 22 with unclear functions) with such domains among the chloroplast proteome of *M. polymorpha* – a species that evidently does not edit its plastid-encoded genes in the same way that *C. cavernum* does. This serves as a reminder that studying diversity often challenges generalizations and it raises the question why dozens of PPR domain-containing proteins are then imported. Considering the versatile functions of these proteins ranging from RNA editing, translation activation, splicing, to even acting as subunits of the ribosomal complex^91–97^, a detailed investigation of each of these proteins will likely reveal novel aspects of chloroplast biology in the liverwort.

In the World Wide Protein Data Bank, 49 protein structures are deposited for the entire bryophyte clade (as of May 2026), juxtaposed to 2684 structures for *A. thaliana* alone. Similarly, while the plastid ribosome synthesizes key components of photosynthesis, chlororibosome structures are available only for *S. oleracea*^98^ and *C. reinhardtii*^99^. This constrains modern evolutionary studies that, in addition to primary sequences work with protein structures^100,101^, are used for the design and optimization of semi-synthetic enzymes^102–104^. Our study deposits structures for 30 bryophyte proteins and 3 rRNAs that constitute two key super complexes of cyanobacterial origin, the large subunit of the chlororibosome and the hetero-oligomeric form I of RuBisCO with its eight small and large subunits. The RAPPL proteome does contain proteins of the small subunit of the chlororibosome (Table S4) and, in combination with the absence of additional assembly factors on the cLSU structure, it suggests the 70S chlororibosome disintegrated during preparation.

Several ribosomal proteins such as L2 to L6, L14, and L16 are among the very few reliable phylogenetic markers across the diversity of life^105^. The main reasons are two-fold: their conserved core structure over 4 billion years of evolution and an almost strict vertical inheritance even in prokaryotes^63,106–108^. The bryophyte’s chlororibosome shares this overall level of structural conservation, but there are noticeable differences concerning the absence of terminal loops in L13, L22 and L23 when compared to the spinach chlororibosome (Fig. 4). The extended loops in L22 and L23 appear eudicot-specific, however, and are hypothesized to form an interface with the thylakoid membrane^59^. Their functional relevance in eudicots remains to be demonstrated, along with proteins that might functionally replace these regions in the liverwort. The absence of the L13 N-terminal extension in the density map – while being present in the annotated gene – implies either a high degree of flexibility, disorder or potentially a post-translational cleavage event. Whatever the reason, this region of the bryophyte’s chlororibosome seems to better resemble its ancestral roots than that of its eudicot relatives on land.

RuBisCO takes a special place in evolution. It is likely the most abundant protein structure on Earth and the vast majority of biomass owes its origin to it, yet it remains one of the least efficient enzymes we know ^109–111^. It also appears resistant to change. Its macromolecular structure has not been noticeably modified across the vast diversity of photosynthetic eukaryotes since its origin as a hetero-oligomeric complex in cyanobacteria some 3 billion years ago^106^ (Fig. 5). The evolution of RuBisCO witnessed the rise of O_2_, a molecule that lowers its efficiency by several orders of magnitude. Despite a strong selection pressure for its active center to exclude oxygen, RuBisCO remained unaffected and, instead, distinct carbon capture mechanisms evolved^110^. When introduced into *E. coli*, not a single nucleotide mutation was observed after 150 chemostat generations for the two genes encoding RuBisCO, despite the prokaryote’s biochemistry mutationally adapting to use CO_2_ as its sole source for synthesizing hexose, pentose and triose sugars instead ^112^. Thus, RuBisCO presents us with an extreme case of an enzyme that is part of a functional module, in which proteins with many interaction partners or those at the center of critical metabolic fluxes, change in sequence significantly slower than their partners^113,114^. The enzyme’s resilience to environmental changes is telling and might explain why its biotechnological optimization remains kind of a Sisyphean task.

The combination of our plastid isolation and the RAPPL protocol, together with cryogenic EM offers a powerful toolset for exploring the structural proteome of large, soluble protein complexes of the plastid stroma. The application of this workflow in a single species illuminates how plastid proteomes have evolved following ancestral genome reduction in the liverwort and structure determination of the ribosome hints at modifications in ribosomal proteins that are eudicot-specific and likely aid in the co-translational insertion of plastid-encoded thylakoid proteins. We conclude that our data and development of a widely applicable and efficient plastid purification protocol can accelerate and expand our understanding of plastid biology across diverse plant lineages.

## Material and methods

### Plant growth conditions

We used ecotype Takaragaire-1 (Tak-1) of the liverwort *Marchantia polymorpha* throughout the study. The wild type gemmae were grown under continuous light (70 μmol m^-2^ s^-1^) on half-strength Gamborg B5 vitamin agar-plates (GVA).

### Plastid isolation

The scaled-down and rapid plastid method applicable to a wide-range of species rests on a protocol published by Eftychis Frangedakis^115^. The plant tissues were lysed using mortar and pestle, in the presence of 5-10 mL isolation buffer (IB; 50 mM HEPES KOH pH 7.5, 0.33 M sorbitol, 1 mM MgCl_2_, 1 mM MnCl_2_, 2 mM EDTA pH 8, 0.5% w/v BSA, 0.5% w/v PVP40, and freshly added 5 mM ascorbate). The lysate was filtered through a pre-soaked (in IB) Mira cloth and centrifuged at 1000 *g*, 10 min. The pellet was resuspended in 200 µL IB and loaded onto a gradient of 70% Percoll^®^ (400 µL at the bottom of the tube) and 30% Percoll^®^ (1 mL overlayed). Density gradient centrifugation was conducted at 4300 *g*, 30 min (in a swing bucket rotor). Broken plastids were gently removed first from the top of the 30% Percoll^®^ layer. Intact plastids were harvested from the 70-30 layer and washed with 10 mL wash buffer (WB; IB without BSA and PVP40) by centrifugation at 3000 *g*, 10 min. The plastid pellet was washed again with 10 mL WB by centrifugation at 2000 *g*, 10 min. The pellet was washed one final time with 1 mL WB by centrifugation at 1000 *g,* 10 min. From the plastid pellets, crude lysate was formed by pulverizing them in presence of liquid nitrogen. Protein isolation buffer (Invitrogen^TM^, Catalogue number FNN0011) was added to the powder and was incubated at 50 °C for 15 min for resuspension. The debris were pelleted at 10,000 *g*, 10 min and the supernatant was used for western blotting.

### Mass spectrometry analysis

The crude plastid extract (obtained as per above) was separated by SDS-PAGE and isolated gel pieces were reduced with 50 µL of 10 mM DTT, alkylated with 50 µL of 50 mM iodoacetamide, and digested with 6 µL (200 ng) trypsin in 100 mM ammonium bicarbonate. The peptides were resolved in 15 µL 0.1% trifluoroacetic acid and used for liquid chromatography. LC-MS data was analyzed using QExactive Plus (Thermo Scientific) connected to an Ultimate 3000 Rapid Separation liquid chromatography system (Thermo Scientific) and equipped with an Aurora Ultimate column (75 µm inner diameter, 25 cm length, 1.7 µm particle size, IonOpticks) as the analytical separation column and an Acclaim PepMap 100 C18 column (75 µm inner diameter, 2 cm length, 3 µm particle size, Thermo Fisher Scientific) as the trap column and with a gradient length of 180 min. The mass spectrometer, operated in positive mode, was coupled to a nano-electrospray ionization source at capillary temperature of 250 °C, source voltage of 1.7 kV. Survey scans were acquired over a mass range of 350–2000 m/z and at a resolution of 140,000. The 10 most intense peptide ions were isolated and fragmented by high-energy collision dissociation (HCD). The peptides were matched against uniport entries for *Marchantia polymorpha* and proteins represented by at least three peptides (of which two unique) and q-value below 0.05 in at least two biological replicates were included in the plastid proteome.

The whole enriched RAPPL sample and 25% of the flow through were precipitated onto magnetic beads (MagReSynAmine, Resyn Biosciences) with 70% acetonitrile. The proteins were washed on the beads with 100% acetonitrile, 70% ethanol and then resuspended in 100 µL of 50 mM NH_4_HCO_3_. The proteins were reduced by the addition of 0.5 M DTT to a final concentration of 10 mM and incubation at 37 °C for 45 min. Then, to alkylate proteins, 2.7 µL of 550 mM IAA was added to a final concentration of 15 mM and samples were incubated at RT in the dark for 30 min. 0.5 µg trypsin was added to each sample for overnight on-bead protein digestion at 37 °C. The resulting peptides were cleaned for mass spectrometry by the STAGE-TIP method using a C18 resin disk (Affinisep).

Samples were then analysed by a nanoElute UHPLC coupled to a timsTOF fleX mass spectrometer (Bruker Daltonics, Bremen, Germany) via a CaptiveSpray ion source. Peptides were separated on a 25 cm reversed-phase C18 column (1.6 µm bead size, 120 Å pore size, 75 µm inner diameter, Ion Optics) with a flow rate of 0.3 µL/min and a solvent gradient from 0-35% B in 60 min. Solvent B was 100% acetonitrile in 0.1% formic acid and solvent A 0.1% formic acid in water. The timsTOF fleX mass spectrometer was operated in DIA-PASEF mode. Mass spectra for MS were recorded between m/z 100 and 1700. Ion mobility resolution was set to 0.85–1.30 V·s/cm over a ramp time of 100 ms. The MS/MS mass range was limited to m/z 475-1000 and ion mobility resolution to 0.85-1.30 V·s/cm to exclude singly changed ions. The estimated cycle time was 0.95 s with 8 cycles using DIA windows of 25 Da. Collisional energy was ramped from 20 eV at 0.60 V·s/cm to 59 eV at 1.60 V·s/cm.

Raw data files from LC-MS/MS analyses were submitted to DIA-NN (version 2.0.2) for protein identification and label-free quantification using *in silico* generated libraries from *Marchantia polymorpha* (trEMBLand Swiss-Prot 2026) and a common laboratory contaminants database from MaxQuant. Carbamidomethyl (C) was set as a fixed modification. Trypsin without proline restriction cleavage option was used, with one allowed miscleavage and peptide length range was set to 7-30 amino acids. MS1 and MS2 mass accuracy was set to 15 ppm and the precursor false discovery rate (FDR) allowed was 0.01 (1%). Match between runs was activated and also RT-dependent cross-run normalization.

### Subcellular localization

Marchantia proteins were localized as previously described^28^. Briefly, full length sequences were amplified from cDNA and fused with citrine on pMpGWB106^116^, through the *Gateway* (Invitrogen) entry vector pDONR221^TM^, by BP and LR reactions set as per the manufacturer’s protocol. The final vectors were transfected via Agrobacterium^117^ and isogenic plant lines were selected on GVA by Hygromycin (10 μg/mL) and Cefotaxime (100 μg/mL). Localization experiments were conducted in protoplasts isolated from 4–7-day old thalli of isogenic transfectant lines. For protoplast isolation, the thalli were incubated with 10 mL of 8% mannitol, 6.3 g/L Gamborg B5 Vitamins (MGV) with 15 mg/mL cellulase and 5 mg/mL Macroenzyme R10. Protoplasts were dislodged and filtered with a 50 µm strainer. After washing twice with MGV (at 300 *g*, 3min) and gently resuspending, the protoplasts were used for mitochondrial staining by MitoTracker™ Red CMXRos according to the manufacturer’s instructions. Protoplasts were imaged using Nikon eclipse Ti imaging platform and collages were prepared in ImageJ (Fiji).

### Comparative organelle proteomics

We clustered 56 Chloroplastida genomes and sorted them into organelle orthogroups as previously described^20^. Briefly, we downloaded whole genome models for diverse Chloroplastida species, clustered them into orthologous protein families (or orthogroups) using OrthoFinder version 2.5.4^118^ and annotated a subset of these orthogroups as plastid orthogroups (POGs) based on the presence of experimentally validated plastid proteins from species where available: *A. thaliana*^36^, maize^35^, rice^35^, *Physcomitrium patens*^34^ and *C. reinhardtii*^32,33^, as well as *M. polymorpha* (current study). Through this approach, more than 99% of ca. 15k experimentally validated plastid proteins were efficiently clustered into ca. 5k POGs (Table S3). The high efficiency is likely explained by the added experimental proteomes and removed non-chloroplastidal species, where homologues are more diverged.

### Ribosome isolation from Marchantia chloroplasts

Isolated plastids were lysed at 4 °C for 30 min with rotation in lysis buffer (2% (v/v) Triton X-100, 50 mM Tris-HCl pH 7.5, 50 mM KCl, 2 mM DTT, 10 mM Mg(OAc)_2_, 2 mM spermidine, 40 U/μL RNase inhibitor murine, 0.1 mM phenylmethylsulfonyl fluoride). Lysate was clarified by centrifugation at 20,800 *g* at 4 °C for 45 min. Purification of the clarified lysate was conducted following the RAPPL protocol. 50 μL poly-*L*-lysine bead slurry (Molecular Cloning Laboratories) was added to lysate supernatant and rotated at 4 °C for 30 min. After placement in a magnetic tube rack, the flowthrough was removed and 250 μL wash buffer (100 mM HEPES pH 7.4, 50 mM KCl, 10 mM Mg(OAc)_2_, 1 mM DTT, 40 U/μL RNase inhibitor murine, 2 mM spermidine) was added. After mixing by pipetting and vortexing, the tube was placed into the magnetic rack and the solution of unbound proteins was removed. The wash was repeated twice (3 in total). 20 μL elution buffer (wash buffer containing 10 mg/mL poly-*D*-glutamic acid) was added to the beads and then incubated at 4 °C with rotation for 30 min.

### Cryogenic EM and TEM imaging

Samples were vitrified on a Leica GP using ANTcryo R1.2/1.3 Au 300 grids. The grids were imaged with a 200 kV Talos Arctica TEM equipped with a Falcon 4i direct electron detector (FINStruct cryo-EM unit, University of Helsinki). Data collection with Thermo Fisher EPU was conducted at 150,000x nominal magnification with a pixel size of 0.92 Å and total electron dose of 39.69 e/Å^2^. In total, 5,652 movies were collected in the .eer format (see Table S1).

### Single-particle processing

Movies were processed in cryoSPARC v4.7^119^. The EER up-sampling factor was set to 1 during movie import. After patch motion correction and patch CTF estimation, pre-processed micrographs were checked and micrographs with an estimated CTF above 5 Å were rejected, leaving 4,934 accepted for further processing. 2,964,872 particles were initially selected using a blob picker and, after inspection, 1,280,024 particles were extracted with a box size of 416 pixels. Extracted particles were run in 2D classification iteratively, using a variable number of 2D classes, 40 online-EM iterations, a batchsize per class of 200, and minimize over per particle scale after the first iteration. 150 2D classes composed of 379,270 particles were used for template picking and 1,309,380 inspected template picked particles were extracted with a box size of 416 pixels. After another round of iterative 2D classification (same parameters as above), 126 classes composed of 364,829 particles were selected. 3D volumes were created using ab-initio reconstruction, with 3 classes, a 260 initial minibatch size, and 900 final minibatch size. All 3 classes were used in a single heterogeneous refinement with a batch size of 2,000 per class and a box size of 416 voxels. 264,465 particles comprised the largest refined class (2.89 Å resolution) which was used for subsequent non-uniform refinement with optimize per-particle defocus, optimize per-group CTF params, and positive EWS correction^120^. The resulting 2.37 Å map was used as the input volume for reference-based motion correction^121^ where movies with wrong frame count were excluded. The resulting 262,814 particles, were used for another non-uniform refinement with the same parameters as above, resulting in a 2.23 Å map (Fig. S4).

Non-ribosomal particles were selected from the blob picked 2D classes for separate analysis using an identical pipeline and similar individual job parameters (Fig. S4). 1,485,133 particles were extracted with a 256 pixel box size. 13 2D classes composed of 471,791 particles were used for ab-initio reconstruction and subsequent heterogeneous refinement using 3 ab-initio volumes. The largest class, composed of 299,945 particles (3.20 Å), was identified as a map of RuBisCO and was used as an input for non-uniform refinement resulting in a 2.52 Å map in C1 symmetry. When D4 symmetry was applied in a separate non-uniform refinement, the result was a 2.21 Å map. After reference-based motion correction and a final non-uniform refinement with D4 symmetry applied, 299,471 particles composed the 2.12 Å map.

Local resolution estimation was conducted for both maps (2.23 Å cLSU and 2.12 Å RuBisCO), with the results analyzed in UCSF ChimeraX v1.9 (Fig. S4,^122^). Both maps were sharpened in DeepEMhancer^123^ (within cryoSPARC) with the 3 available models used for both maps and manually compared in USCF ChimeraX, with the reported maps sharpened using the wideTarget model for RuBisCO and highRes model for the LSU maps respectively.

### Modeling and validation

Protein sequences were obtained through BLASTP starting from 5MLC *S. oleracea* 50S chlororibosome structure and 7JN4 *C. reinhardtii* RuBisCO looking for *M. polymorpha* sequences^124^. The following GenBank accession numbers were used for 50S sequences: BBN04705.1, PTQ33360.1, OAE33939.1, P12142.1, OAE31853.1, OAE19889.1, P06392.3, P12196.3, OAE31933.1, OAE33704.1, PTQ42878.1, PTQ48266.1, BDD77304.1, P06388.1, P06387.1, P06385.3, PTQ47902.1, PTQ36989.1, PTQ46014.1, YP_009479654.1, OAE34979.1, YP_009479653.1, OAE23259.1, PTQ48626.1, OAE20865.1, APT68014.1, PTQ48664.1, P06378.1, and RuBisCO sequences: ALV87782.1, PTQ31246.1^53,54,56–58^. BLASTP sequences were run through AlphaFold3 on the publicly-accessible web server and then fit to the previously mentioned PDB structures using the Matchmaker tool in UCSF ChimeraX. They were subsequently fit to their respective density maps using the Fit in Map tool in UCSF ChimeraX. Models were fit residue by residue using ISOLDE in ChimeraX and regions of the sequences with no corresponding density were deleted from the models (Table S6)^125^.

Nucleotide sequences were obtained through BLASTN starting from the PDB ID 5MLC spinach chlororibosome structure with the following GenBank accession numbers: NC_042505.1, KT356925.1, KT356925.1^55,59^. The 5MLC structure was used as a starting point for rRNA structures. Individual point mutations and deletions were made in UCSF ChimeraX using swapnt and delete commands. Insertions were made with the Build Structure tool in UCSF ChimeraX and Coot v0.9.8.96, which was used to re-order and change chain IDs^126^. Models were fit residue by residue using ISOLDE and regions of the sequences with no corresponding density were deleted from the models (Table S6).

Ions were fit to the cLSU structure using the isolde add ligand command where a strong signal with appropriate coordination was present in the sharpened density map. Specific ion assignment was based on coordination distances and interacting atoms. Coordination was confirmed with water molecules using the un-sharpened map and the isolde add water command. Spermidine was added based on unmodeled density in the sharpened map with the Build Structure tool. rRNA modifications were added based on unmodeled density in the sharpened map with the Build Structure tool. All ligands and modifications were fit with refinement in ISOLDE. Validation was conducted during modeling in ISOLDE and by the wwPDB OneDep validation servers.

## Data availability

Structures were deposited to the Worldwide Protein Data Bank through the wwPDB OneDep system. The cLSU structure was given the following accession codes: PDB ID 31DT, EMD-58329. The RuBisCO structure was given the following accession codes: PDB ID 31EJ, EMD-58338. Micrographs are deposited through EMPIAR (deposition ID: 47488363, accession code: EMPIAR-13780) and linked to both Electron Microscopy Data Bank accessions. Supplementary figures are available with this submission. Additional data are available on Zenedo (https://zenodo.org/records/21338199), which includes, but is not limited to, figure source data, a supplementary movie for *M. polymorpha* chloroplast ribosome LSU, supplementary tables and in-house scripts.

## Author contributions

PKR: Conceptualization, Experimental design, Methodology, Investigation, Data curation, Formal analysis, Validation and Visualization, Writing - original draft, review and editing. CM: Experimental design, Methodology, Investigation, Data curation, Formal analysis, Visualization, Writing - original draft, review and editing MLQ: Investigation; Data curation and Validation. SOK: Experimental design, Methodology; Funding and Resource acquisition; Writing - review and editing; TAN: Formal analysis; BJB: Supervision; Experimental design; Funding and Resource acquisition; Writing - review and editing; SJB: Supervision; Experimental design; Funding and Resource acquisition Writing - review and editing. SBG: Conceptualization; Project administration; Supervision; Experimental design; Funding and Resource acquisition; Data visualization; Writing - review and editing.

## Acknowledgments

We acknowledge support from the high-performance computing cluster (HILBERT) at the Heinrich Heine University Düsseldorf and thank Daniel Wasim Djamriani, Carolina Garcia Garcia and Margarete Stracke for their help. We also thank Dr. Anja Stefanski and Molecular Proteomics Laboratory at HHU-Düsseldorf for conducting the mass-spectrometry. PKR and MLQ are grateful to Prof. William F. Martin for providing financial support. Mass spectrometry-based proteomic analyses of RAPPL sample preparations were performed by the Proteomics Core Facility, Department of Immunology, University of Oslo/Oslo University Hospital, which is supported by the Core Facilities program of the South-Eastern Norway Regional Health Authority. This core facility is also a member of the National Network of Advanced Proteomics Infrastructure (NAPI), which is funded by the Research Council of Norway INFRASTRUKTUR-program (project number: 295910). We thank Kiran L.L. Ahmad and Tuomas Niemi-Aro (University of Helsinki) for technical assistance in cryo-EM. The facilities and expertise of the HiLIFE cryo-EM unit at the University of Helsinki, a member of Instruct-ERIC Centre Finland, FINStruct, and Biocenter Finland are gratefully acknowledged. We acknowledge computational resources from CSC – IT Center for Science, Finland. Molecular graphics and analyses performed with USCF ChimeraX, developed by the Resource for Biocomputing, Visualization, and Informatics at the University of California, San Francisco, with support from National Institutes of Health R01-GM129325 and the Office of Cyber Infrastructure and Computational Biology, National Institute of Allergy and Infectious Diseases.

