## Supplementary Figures 1-7 for "Rapid plastid isolation reveals the chloroplast proteome and structures of the chlororibosome large subunit and RuBisCO in *Marchantia polymorpha*"

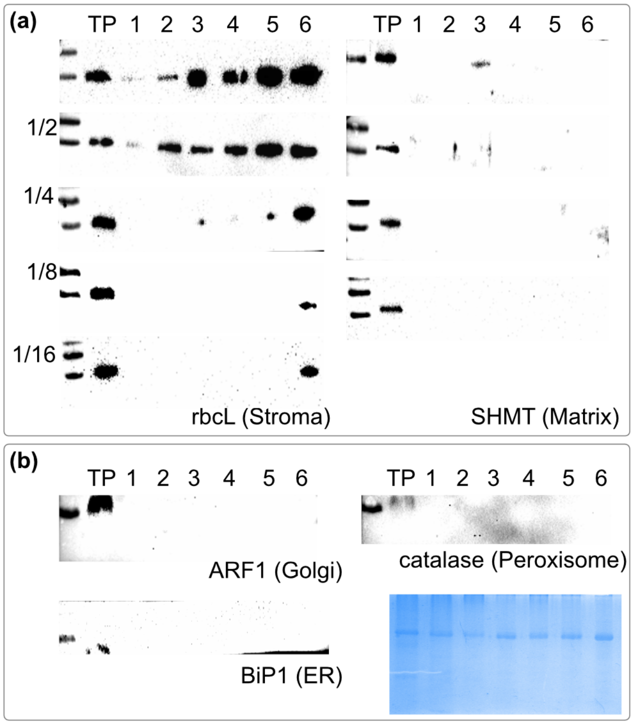


**Figure S1: Enrichment of plastids from *Marchantia*.** **(a)** Immunoblotting from total protein (TP) and chloroplast protein from six independent chloroplast preparations from ca. 10 g (Prep 1-2), 20 g (Prep 3-4) and 30 g (Prep 5-6) thalli against rbcL (stroma marker) and SHMT (mitochondrial marker). The plastid proteins serially diluted from 1/2 to 1/16 underscore the enrichment of the isolated plastids. **(b)** Immunoblotting against marker proteins for the Golgi, ER, peroxisome and Coomassie-staining showing even sample loading. A prestained protein ladder is loaded in the first well of each gel.

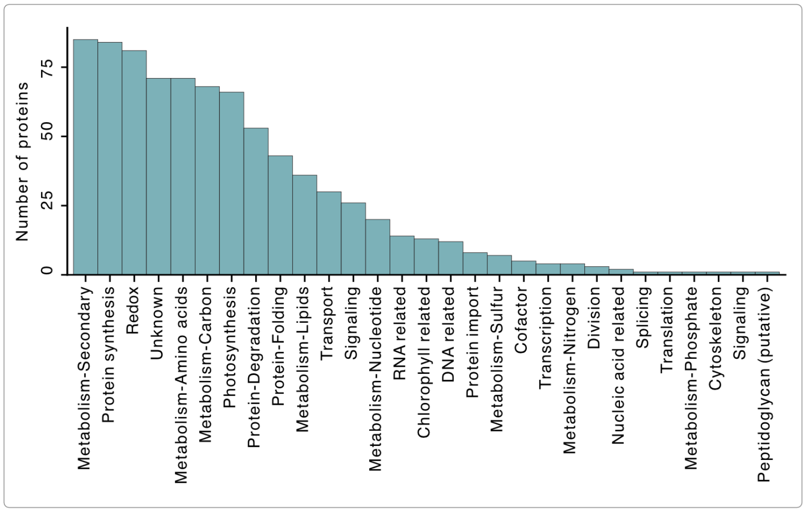


**Figure S2: Cyanobacterial homologues of *M. polymorpha* plastid proteins.** Functions of 812 *M. polymorpha* plastid proteins with homologues in a database derived from 56 cyanobacteria. Another 527 *M. polymorpha* plastid proteins lack homologues in Cyanobacteria.

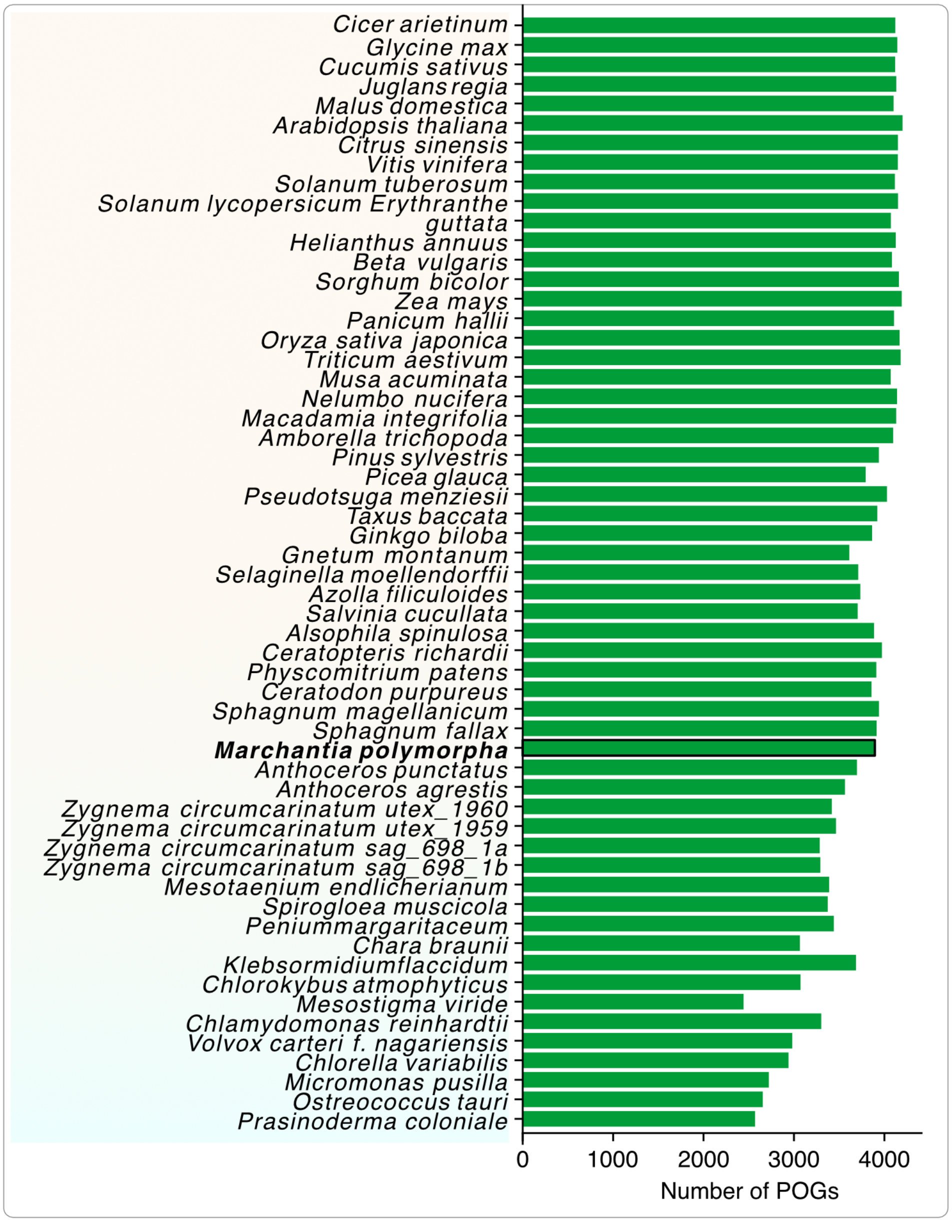


**Figure S3: Number of plastid localized orthogroups per species across 57 diverse chloroplastida species.**

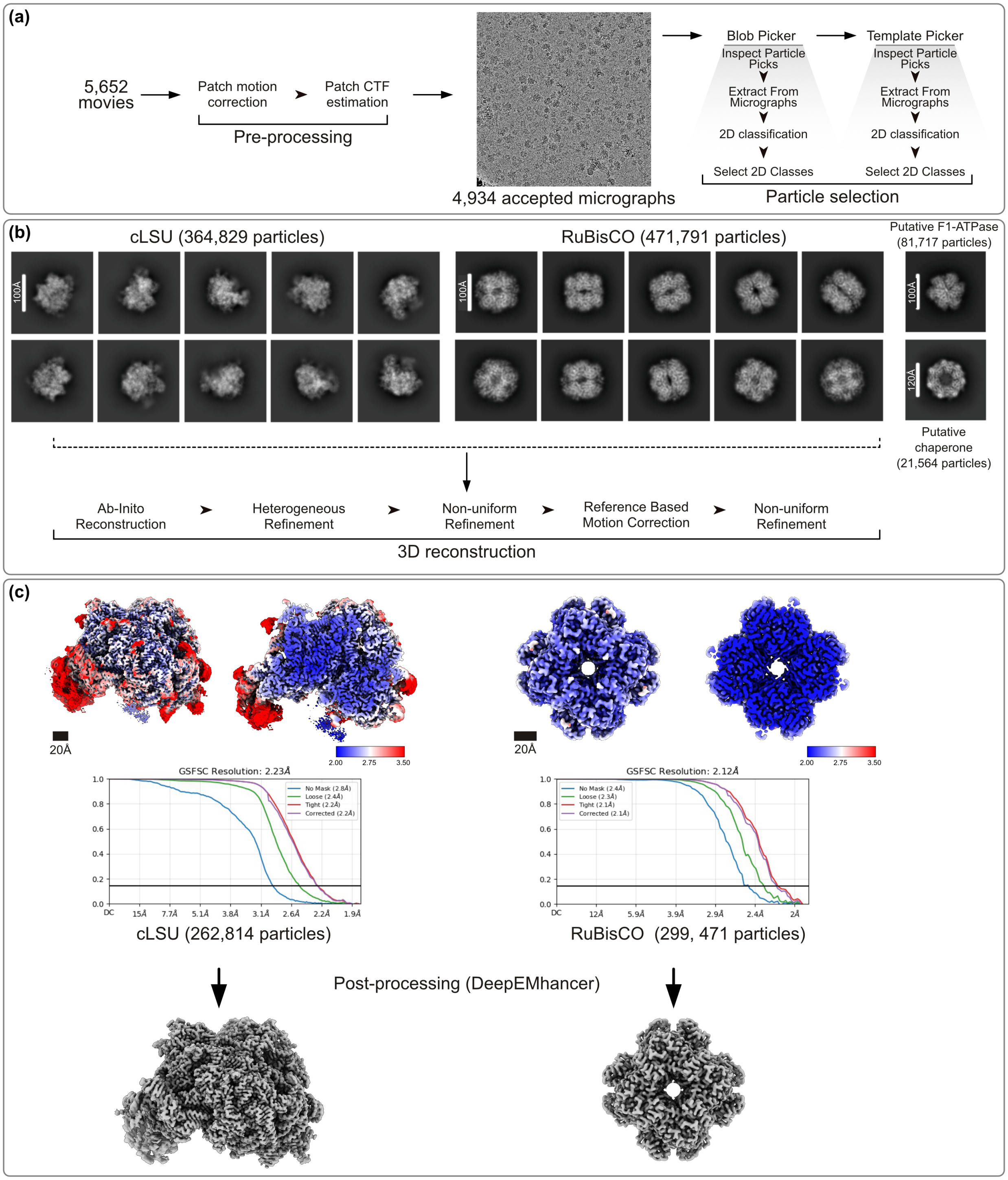


**Figure S4: CryoSPARC processing workflow and exemplary results. (a)** 5,652 cryoEM movies were preprocessed, resulting in 4,934 micrographs meeting the quality thresholds. Particles were selected from these micrographs via Blob Picker and Template Pickers, with a micrograph shown. **(b)** The workflow resulted in ca. 300-400K particles for chloroplast large subunit (cLSU) and RuBisCO, alongside <100K particles of putative F1-ATPase and Chaperone, with example 2D class averages shown. From the cLSU and RuBisCO particles, 3D images were reconstructed at high resolution, with the local resolution maps (left: surface, right: central cross section) and FSC curves shown. **(c)** Density maps were sharpened via DeepEMhancer, and the resulting sharpened maps are shown.

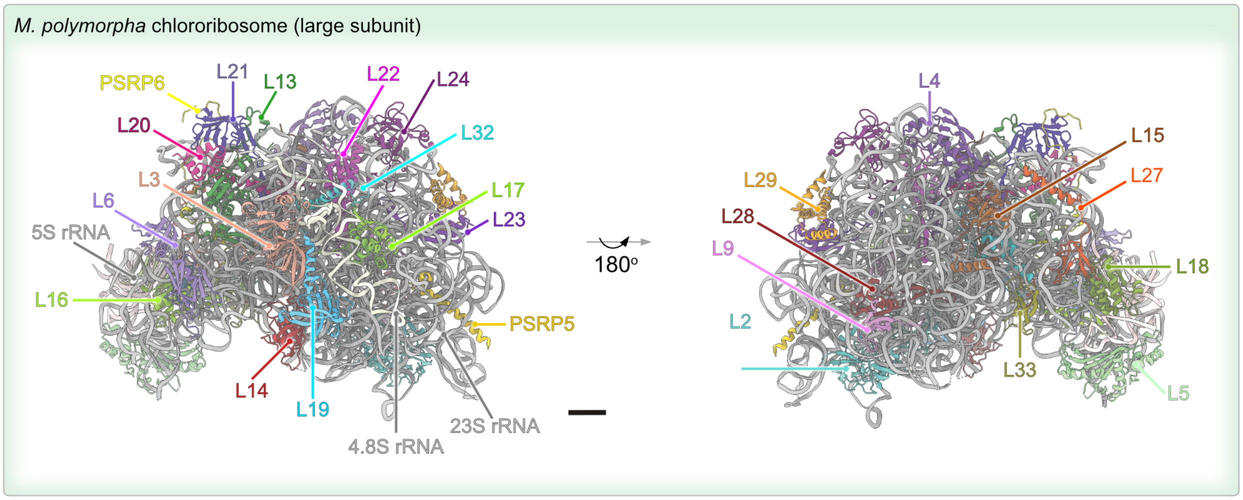


**Figure S5: Structure of Marchantia chlororibosomal large subunit.** Atomic model of *M. polymorpha* cLSU, as ribbon representation, colored according to individual RPs and rRNAs**.**

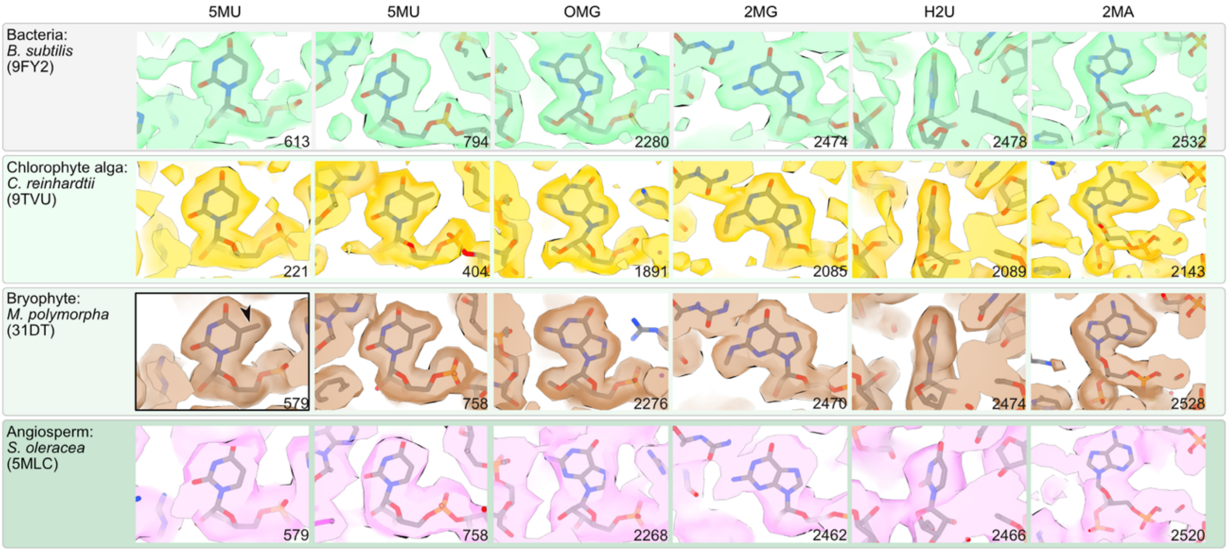


**Figure S6: rRNA modification from chlororibosomes across species.** Modifications in the 23S rRNA of chlororibosomes with the bases labeled on the bottom right in each image. A modification likely unique to *M. polymorpha* is indicated by a black border and an arrow within the image. (5MU = 5-methyluridine, OMG = 2′-O-methylguanosine, 2MG = 2N-methylguanosine, H2U = 5,6-dihydrouridine, 2MA = 2-methyladenosine).

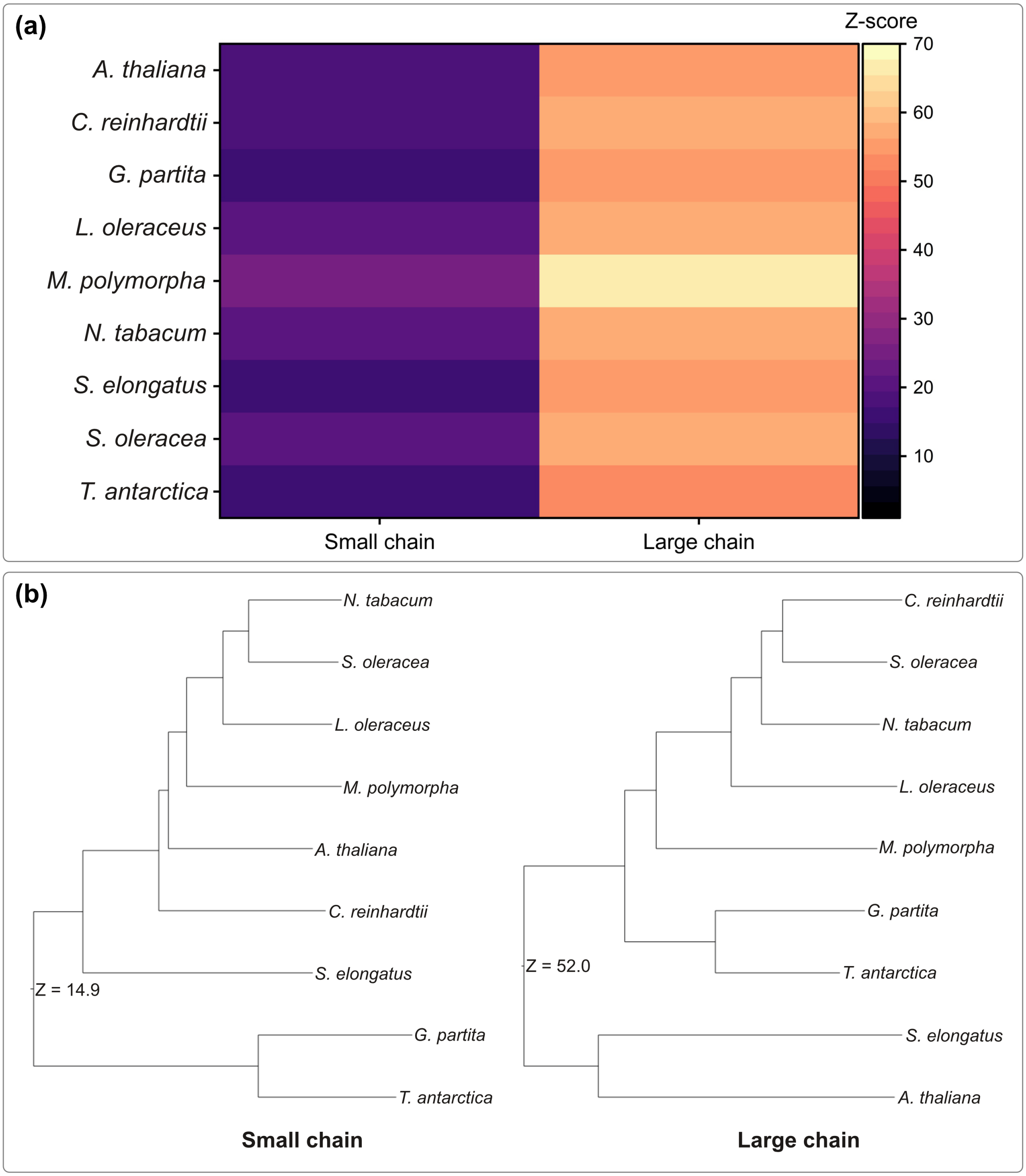


**Figure S7: Structural comparison of RuBisCO across species.** DALI Z-score across nine species represented as a heatmap **(a)** and cladograms **(b)**. (*A. thaliana* = *Arabidopsis thaliana, C. reinhardtii* = *Chlamydomonas reinhardtii, G. partita* = *Glaucocystis partita, L. oleraceus* = *Lophomonas oleraceus, M. polymorpha* = *Marchantia polymorpha, N. tabacum* = *Nicotiana tabacum, S. elongatus* = *Synechococcus elongatus, S. oleracea* = *Spinacia oleracea, T. antarctica* = *Trebouxia antarctica*). Compared structures and their respective PDB IDs are referred to in Fig. 5.
