## Supplementary material for "Rapid plastid isolation reveals the chloroplast proteome and structures of the chlororibosome large subunit and RuBisCO in *Marchantia polymorpha*": Table S6

**Table 1. Cryo-EM data collection, refinement and validation statistics**

|  | **50S** | **RuBisCO** |
| --- | --- | --- |
| **Data collection and processing** |  |  |
| **Magnification** | 150,000 | 150,000 |
| **Voltage (kV)** | 200 | 200 |
| **Electron exposure (e–/Å^2^)** | 39.69 | 39.69 |
| **Defocus range settings (μm)** | -0.6 to -2.0 | -0.6 to -2.0 |
| **Pixel size (Å)** | 0.92 | 0.92 |
| **Symmetry imposed** | C1 | D4 |
| **Micrographs (no.)** | 4,934 | 4,934 |
| **Final particle images (no.)** | 262,814 | 294,936 |
| **Map resolution (Å)**  **FSC threshold** | 2.23  0.143 | 2.12  0.143 |
| **Map resolution range (Å)** | 999 – 1.84 | 999 – 1.84 |
| **Refinement** |  |  |
| **Model composition**  **Non-hydrogen atoms**  **Protein residues** | 88,889  3,307 | 36,408  4,608 |
| **Small chain**  **Large chain**  **L2** | -  -  2-234, 245-276 | 132-253  12-465  - |
| **L3** | 82-301 | - |
| **L4** | 68-278 | - |
| **L5**  **L6**  **L9**  **L13**  **L14**  **L15**  **L16**  **L17**  **L18**  **L19**  **L20**  **L21**  **L22**  **L23**  **L24**  **L27**  **L28**  **L29**  **L32**  **L33**  **L34**  **L35**  **L36**  **PSRP5**  **PSRP6**  **Nucleotide residues**  **4.8S**  **5S**  **23S** | 83-227, 233-265  49-226  52-101  46-224  1-122  76-260  1-135  93-208  65-186  119-237  2-116  2-116  8-119  1-90  48-182  63-171  53-131  57-151  2-50  5-65  106-163  77-145  1-37  143-189  38-93  2,866  1-103  3-121  1-896, 901-1093, 1098-1101, 1109-1119, 1126-1512, 1521-1854, 1869-1891, 1895-1919, 1966-1971, 1986-2124, 2198-2810 | -  -  -  -  -  -  -  -  -  -  -  -  -  -  -  -  -  -  -  -  -  -  -  -  -  -  -  -  - |
| **Ligands** | H_2_O, K^+^, Mg^2+^, Na^+^, Zn^2+^, Spermidine | - |
| **Validation** |  |  |
| **R.m.s. deviations**  **Bond lengths (Å)**  **Bond angles (°)** | 0.00  0.18 | 0.00  0.22 |
| **Clashscore** | 0.08 | 1.52 |
| **Rotamer outliers (%)** | 0.22 | 0.00 |
| **Ramachandran plot** |  |  |
| **Favored (%)** | 97.43 | 96.68 |
| **Allowed (%)** | 2.48 | 3.32 |
| **Disallowed (%)** | 0.10 | 0.00 |
